# Exploring the prediction of emotional valence and pharmacologic effect across fMRI studies of antidepressants

**DOI:** 10.1101/382408

**Authors:** Daniel Barron, Mehraveh Salehi, Michael Browning, Catherine J Harmer, R. Todd Constable, Eugene Duff

**Author notes:** Corresponding Author: Daniel S. Barron 300, George Suite #901, New Haven, CT 06510.

## Abstract

**Background:** Clinically approved antidepressants modulate the brain’s emotional valence circuits, suggesting that the response of these circuits could serve as a biomarker for screening candidate antidepressant drugs. However, it is necessary that these modulations can be reliably detected. Here, we apply a cross-validated predictive model to classify emotional valence and pharmacologic effect across eleven task-based fMRI datasets (n=306) exploring the effect of antidepressant administration on emotional face processing.

**Methods:** We created subject-level contrast of parameter estimates of the emotional faces task and used the Shen whole-brain parcellation scheme to define 268 subject-level features that trained a cross-validated gradient-boosting machine protocol to classify emotional valence (fearful vs happy face visual conditions) and pharmacologic effect (drug vs placebo administration) within and across studies.

**Results:** We found patterns of brain activity that classify emotional valence with a statistically significant level of accuracy (70% across-all-subjects; range from 50-87% across-study). Our classifier failed to consistently discriminate drug from placebo. Subject population (healthy or unhealthy), treatment group (drug or placebo), and drug administration protocol (dose and duration) affected this accuracy with similar populations better predicting one another.

**Conclusions:** We found limited evidence that antidepressants modulated brain response in a consistent manner, however found a consistent signature for emotional valence. Variable functional patterns across studies suggest that predictive modeling can inform biomarker development in mental health and in pharmacotherapy development. Our results suggest that case-controlled designs and more standardized protocols are required for functional imaging to provide robust biomarkers for drug development.

## 1. INTRODUCTION

Psychiatric drug development is difficult, expensive, and beset by a high failure rate. The slow onset, unclear biological markers, and variable clinical efficacy even of approved psychiatric drugs makes the potential efficacy of candidate drugs difficult to measure and has led many pharmaceutical companies to withdraw from drug development (1; 2). Biomarkers that capture how effective drugs modulate the brain’s functional anatomy could prioritize candidate compounds for large clinical trials, thus improving the productivity and cost-effectiveness of drug development.

Clinically approved antidepressants modulate the brain’s emotional valence circuits, suggesting that the response of these cicruits could serve as a biomarker for screening candidate antidepressant drugs. The emotional faces task has been particularly useful in eliciting the emotional valence circuit (3; 4). In this task, a subject is instructed to view a human actors’ face and determine the gender of or the emotion expressed. Independent studies have shown that emotional valence networks engaged by this task are affected by antidepressant administration (5). The applicability of these studies to screen for potential antidepressant compounds rests on the ability of the emotional faces task to engage a spatially consistent emotional valence network across populations, specifically the aspect of this network that is affected by antidepressant administration. This applicability may be explored by assessing 2 contrasts: an emotional valence contrast (i.e. is there a consistent difference in activity when positive and negative faces are displayed?) and a pharmacologic contrast (is there a consistent difference when antidepressants are compared to placebos?).

A second advantage of the emotional valence contrast described above is that it can be constructed in either a within or between-subject manner. Duff et al (2015) have previously successfully developed a cross-validated machine learning protocol which was able to predict pharmacologic class in analgesic studies within pain stimulation tasks. However, the analgesia literature tends to use within subject designs whereas the antidepressant literature uses between subject designs. The emotional valence contrast is therefore useful as a means of directly comparing classifier performance of within vs. between subject contrasts on the same dataset.

Here, we apply a machine-learning classifier to a large set of studies of antidepressant effects on brain responses during an emotional faces tasks. We explore the consistency of the emotional valance effect considered both within and between-subjects and the between-subject pharmacologic effect. Because these studies use protocols with considerable variability in scanners, experimental tasks and patient cohorts, we further aim to explore the effect of protocol variability on signature generalizability. To accomplish this, we exploit a dimensionality reduction step(6) to reduce voxel-wise data to functionally homogenous parcels defined in an independent dataset by an unsupervised algorithm (7). We then apply the gradient boosted machine (GBM) classifier to predict emotional valence (fearful vs happy face presentation) and pharmacologic class (antidepressant versus placebo), to test whether a consistent, cross-study signature may be identified, and to understand which study protocols generate a more generalizable signature.

## 2. METHODS AND MATERIALS

For each of eleven datasets, subject-level contrast of parameter estimates of the emotional faces task were created and divided into 268 regions using the Shen whole-brain parcellation scheme. Each region was used as a feature within a cross-validated gradient-boosting machine protocol that classified emotional valence and pharmacologic effect within and across studies. Feature weightings were then mapped onto the brain to allow anatomic localization and visualization.

### 2.1 Datasets

Eleven independent datasets from eight task-based fMRI studies of the effect of antidepressant administration on emotional face processing were available for analysis, representing 306 subjects (See Table 1 for key features of the dataset; NB: the number of subjects per study differs from the original publications, reflecting that some data could not be located for inclusion in our study and that one study (Warren) has recruited more participants since the time of our study). These studies were all performed in the Harmer lab from 2006-2015 and made use of healthy subjects (H) without previous history of mental illness and subjects selected based on the presence of symptoms consistent with a disorder (i.e. Major Depressive Disorder) or symptom (i.e. neuroticism or dysphoria). In these studies, the Beck Depression Inventory and the Eysenck Personality Questionnaire, neuroticism dimension were used to assess these symptoms. Although specific aspects of the study varied (e.g. antidepressant dose and duration), all versions investigated group differences in whole-brain BOLD response when subjects viewed happy and fearful faces. In this study, we selected only happy and fearful emotional face presentation, as these were the most consistently used emotions in our available dataset. Individual studies each obtained ethical approval from the local ethics committee.

**Table 1.** Summary of Included Datasets.

| Study | Year | Scanner | TR<br>(sec) | Presentation | Task Presentation |  | Emotional Face<br>Stimulus Length<br>(msec) | Drug Administration |  | Clinical<br>Condition | Patient Population* |  | Total<br>(n) |
| --- | --- | --- | --- | --- | --- | --- | --- | --- | --- | --- | --- | --- | --- |
|  |  |  |  |  | Instructions | Acquisition<br>Length<br>(TRs) |  | Drug | Protocol |  | Drug<br>(n) | Placebo<br>(n) |  |
| Harmer | 2004 | 1.5T Siemens<br>Sonata | 3 | Masked | Identify<br>Gender | 190 | 17 (followed by<br>167 neutral face) | Citalopram<br>(SSRI) | 20mg/day,<br>7 days | Healthy | 9 | 8 | 17 |
| Murphy | 2009 | 1.5T Siemens<br>Sonata | 3 | Unmasked | Identify<br>Gender | 330 | 200 (unmasked) | Citalopram<br>(SSRI) | 20mg,<br>single dose | Healthy | 13 | 11 | 24 |
| Rawlings | 2010 | 1.5T Siemens<br>Sonata | 2 | Unmasked | Identify<br>Gender | 250 | 100 | Mirtazapine<br>(NaSSA) | 15mg,<br>single dose | Healthy | 14 | 14 | 28 |
| KumarA | In prep | 3T Siemens | 3 | Unmasked | Identify<br>Gender | 250 | 100 | Citalopram<br>(SSRI) | 20mg/day,<br>7 days | Healthy | 16 | 15 | 31 |
| KumarB | In prep | 3T Siemens | 3 | Unmasked | Identify<br>Gender | 250 | 100 | Citalopram<br>(SSRI) | 20mg/day,<br>7 days | Healthy | 17 | 13 | 30 |
| Warren | In prep | 3T Siemens | 3 | Unmasked | Identify<br>Gender | 270 | 500 | Escitalopram<br>(SSRI) | 20mg/day,<br>7 days | Low<br>Neurotic | 19 | 12 | 31 |
| Warren | In prep | 3T Siemens | 3 | Unmasked | Identify<br>Gender | 270 | 500 | Escitalopram<br>(SSRI) | 20mg/day,<br>7 days | High<br>Neurotic | 14 | 15 | 29 |
| DiSimplicio | 2013 | 3T Siemens | 3 | Unmasked | Identify<br>Gender | 250 | 500 | Citalopram<br>(SSRI) | 20mg/day,<br>7 days | High<br>Neurotic | 14 | 7 | 21 |
| KumarA | In prep | 3T Siemens | 3 | Unmasked | Identify<br>Gender | 250 | 100 | Citalopram<br>(SSRI) | 20mg/day,<br>7 days | Dysphoric | 9 | 18 | 27 |
| KumarB | In prep | 3T Siemens | 3 | Unmasked | Identify<br>Gender | 250 | 100 | Citalopram<br>(SSRI) | 20mg/day,<br>7 days | Dysphoric | 16 | 14 | 30 |
| Godlewska | 2012 | 3T Siemens | 2 | Unmasked | Identify<br>Gender | 250 | 100 | Escitalopram<br>(SSRI) | 10mg/day,<br>7 days | MDD | 19 | 19 | 38 |
| TOTALS |  |  |  |  |  |  |  |  |  |  | 160 | 146 | 306 |

\* The number of subjects per study differs from the original publications. This reflects that some data were unable to be located for inclusion in our study and that one study (Warren) has recruited more participants since the time of our study.

### 2.2 MRI Processing

Standard preprocessing and mapping analysis were employed using tools from FMRIB’s Software Library (FSL) package (http://fsl.fmrib.ox.ac.uk/fsl/fslwiki/). The FSL FMRI Expert Analysis Tool (FEAT) was used for general linear modeling (GLM) (8). Subject-level contrast of parameter estimate (COPE) maps for each contrast (e.g. happy versus fixation) were produced in native patient space. These COPE maps were used in subsequent classification analyses, as described below. See Supplementary Methods for more details and Figure 1 for an illustration of the analysis pipeline.

**Figure 1.**
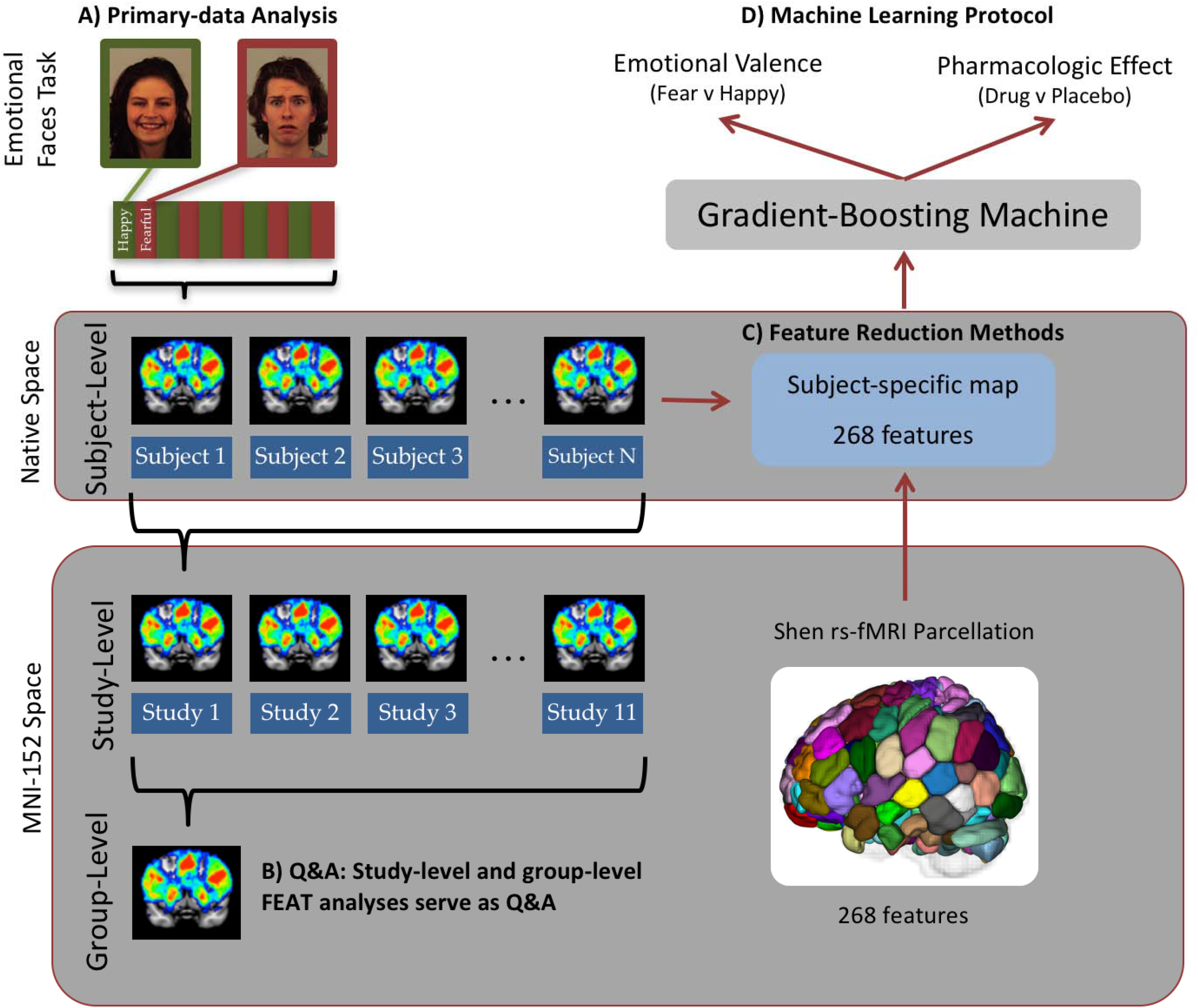
Protocol Summary. Primary-data analysis (A) was performed at the subject level to model task effects. Study and group-level analyses took place in MNI152 space and served as a QA step (B, see Methods). Feature reduction (C) took place in native subject space to maximize registration accuracy. The contrast of parameter estimates (COPE, see Methods) were used as features in the machine learning protocol (D). Illustrative faces in (A) are from Karolinska Institutet’s publically available database (24).

### 2.3 Machine Learning Method

Cognitive models of depression suggest that patients process negative relative to positive stimuli differently from non patients, and that these cognitive processes are causative in the illness. Therefore a contrast looking at the emotional processing circuit activation to negative vs. positive faces may be able to identify illness specific signatures and how the brain’s emotional circuits change in response to treatment. We chose a forced-choice gradient boosting machine (GBM) for classification due to its robustness to outliers and its ability to map features back into anatomical brain space (9).

Predictive analyses are prone to overfitting when the number of features far outweighs the number of subjects(6). Given our available dataset of 306 subjects, we had to reduce the number of features from voxels (~900,000 in 2mm isotropic space). To this end, we selected the Shen 268-node resting-state fMRI atlas, defined by a group-wise spectral clustering algorithm applied to an independent dataset consisting of 45 subjects (7; 10). We transformed the Shen atlas from MNI-152 space into native patient space using linear and nonlinear FSL transforms (8) and used the average COPE values within each parcel to produce 268 features per subject for the classifier.

We trained 2 overall types of classifiers:

1. Emotional Valence Classifier. This analysis determined whether and where a signal for emotional valence was consistent enough to discriminate fear from happy face visual conditions. We assessed the performance of the emotional valence classifier with two different types of feature inputs to determine the impact of inter-subject variability and task variability. The first subtracted fear and happy responses within-subject, to account for average differences in visual responses across subjects (i.e. the classifier compared the FvH COPE contrast image to the HvF COPE contrast image). The second compared fear versus fixation COPE files and happy versus fixation COPEs and accounted for across-study differences in task, without being able to minimize individual subject variability in the visual response. Duff et al (11) were able to minimize intersubject variability through within-subject contrasts wherein each subject received a placebo and drug condition, thus allowing pharmacologic effect to be isolated from variability due to individual differences and/or task. Because the pharmacologic effect in our studies was necessarily between subjects, we used the valence contrast to compare the performance of a within vs. between subject classifier as the structure of the task allowed us to do this.
2. Pharmacologic Effect Classifier. This analysis used contrasts between Fear and Happy conditions to discriminate patients with drug or placebo protocols within and across studies (i.e. the classifier compared the FvH_drug_ COPE contrast image to the FvH_placebo_ contrast image). We further observed whether this signal was consistent across antidepressant dose, frequency, and duration.

We tailored the predictive pipeline and cross validation strategy based on the level of the classification performed; these details (including an explanatory figure) are available in Supplementary Methods. In brief: (i) within-study classification: when subjects within one study were considered (i.e. to determine the reliability of effects within individual studies), the classifier was trained on all but two subjects (balanced for classification group) and then tested on those held out subjects in an iterative fashion, until the classifier was tested on all subjects; (ii) across-study classification: when subjects across two studies were considered (i.e. to assess the similarity of trained classifiers across individual studies), the classifier was trained on one study and tested on the other, (iii) across-all-subjects classification: when subjects across all studies were considered, to determine our ability to build classifiers that generalize across subjects, the classifier was trained on all but two subjects and then tested on those held out subjects in an iterative fashion, until the classifier was tested on all subjects; (iv) across-all-studies classification: when subjects across all studies were considered (i.e. to assess how a classifier trained on all studies performed on a held out study), the classifier was trained on all studies except one and tested on that held out study.

## 3. RESULTS

### 3.1 Emotional Valence: Happy from Fearful Face Classification

We first describe results associated with discrimination of parameter images subtracting fear and happy responses within-subject, which addresses individual variability in visual responses.

i. Within-study classifications (i.e. classifiers trained and tested on different subjects of same study) provide insight into those studies with the most discriminative signals. The five datasets from the Harmer, Murphy and Kumar studies provide the best performance (67-87% accuracy p<0.001; uncorrected for multiple comparison as each study was considered separately. See Table 2, p-values for each accuracy score are shown in the Supplementary Materials Figure 1). These five datasets represented participants who were healthy or showed dysphoric traits. Accuracies were not better than chance for the Rawlings study of healthy participants, for the Warren and Disimplicio studies of participants with low or high neurotic traits, or for the Godlewska study of participants diagnosed with MDD.
ii. Across-study classifications (i.e. classifiers trained on one study and tested on separate studies) show the ability of classifiers trained on one study to discriminate other studies. We found that results varied considerably depending on the study population evaluated. Classifiers trained and tested on studies of healthy, dysphoric, or major depressive disorder (MDD) performed notably better than studies of low or high neurotic trait (p<0.005, (α/10) Bonferroni correction for multiple comparisons given 10 classifications per study). We were unable to find a consistent discriminative signal within studies of neurotic subjects. Each train-test dyad may be referenced in Figure 2. Results when the classifier was trained on all studies or all healthy studies and tested on a held-out study may be referenced in Supplementary Materials Figure 1.
iii. Across-all-subjects classification achieved an average accuracy of 69% (p<0.001) in held out data. Classifiers trained and tested on only healthy subjects achieved an average accuracy of 66% (significant at p<0.005 level, no test for multiple comparinsons.). (iv) Across-all-studies classifications are presented in Supplementary Results.

**Figure 2.**
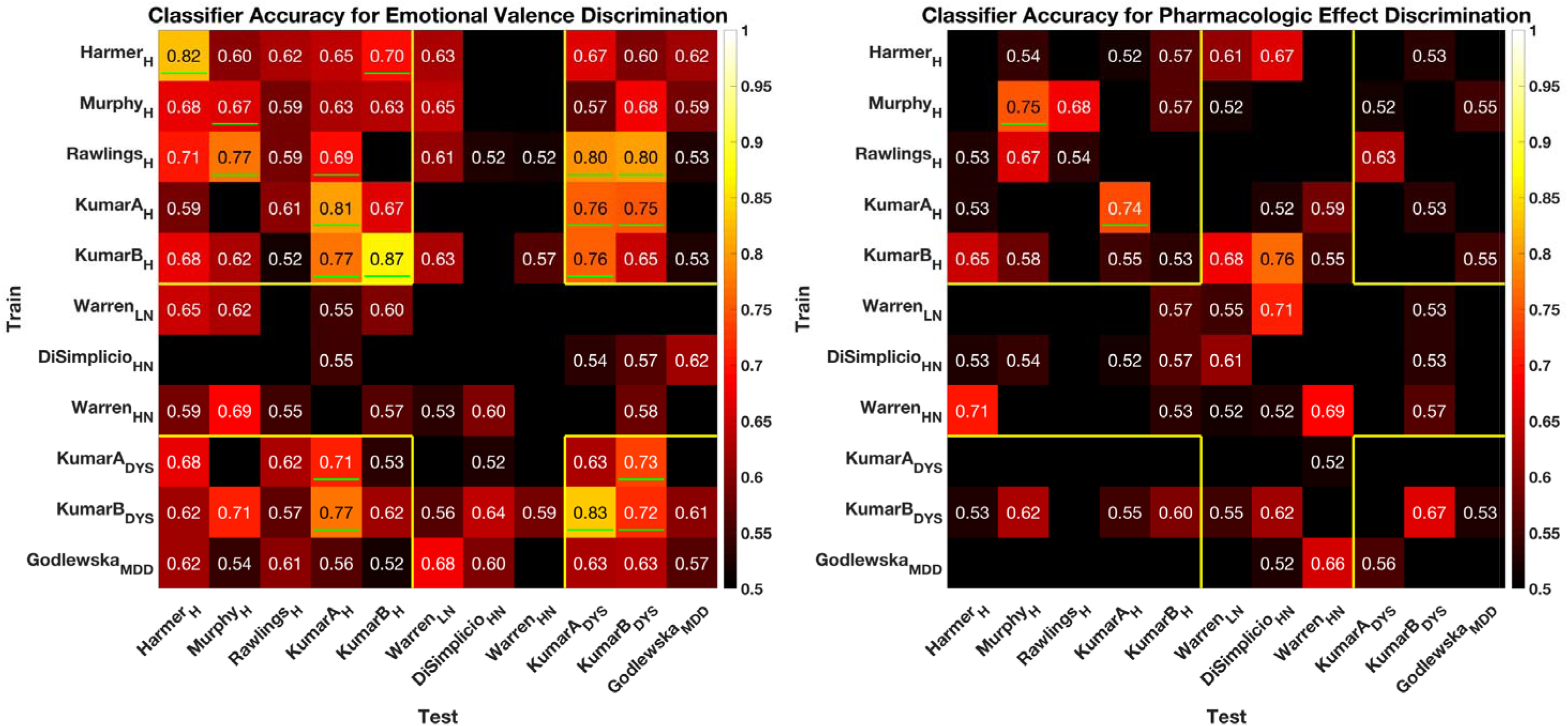
Accuracies for the emotional valence (left) and pharmacologic effect (right) classification. Studies are organized on a clinical spectrum, from healthy (H), to low neurotic (LN), to high neurotic (HN), to dysphoric (DYS), to major depressive disorder (MDD). Green lines indicate significance at respective level: (i) within study classification: no correction for multiple comparisons; (ii) across-study: p<(0.05/10) Bonferroni correction for multiple comparisons; (iii) across all-subjects: no correction for multiple comparisons; (iv) across-all-studies p<(0.05/10) Bonferroni correction for multiple comparisons. Accuracies based on a bimodal distribution test, numerical p-values are shown in the Supplementary Materials. Yellow lines are illustrate groups with higher shared accuracies. Shown below are results for the within-study classification (diagonal) and across-study classification (off-diagonal). Results for the across-all-subjects classification may be referenced in Table 2 (final row) and Supplementary Figure 2. Results for the across-all-studies classification may be referenced in Supplementary Figure 2.

**Table 2.**
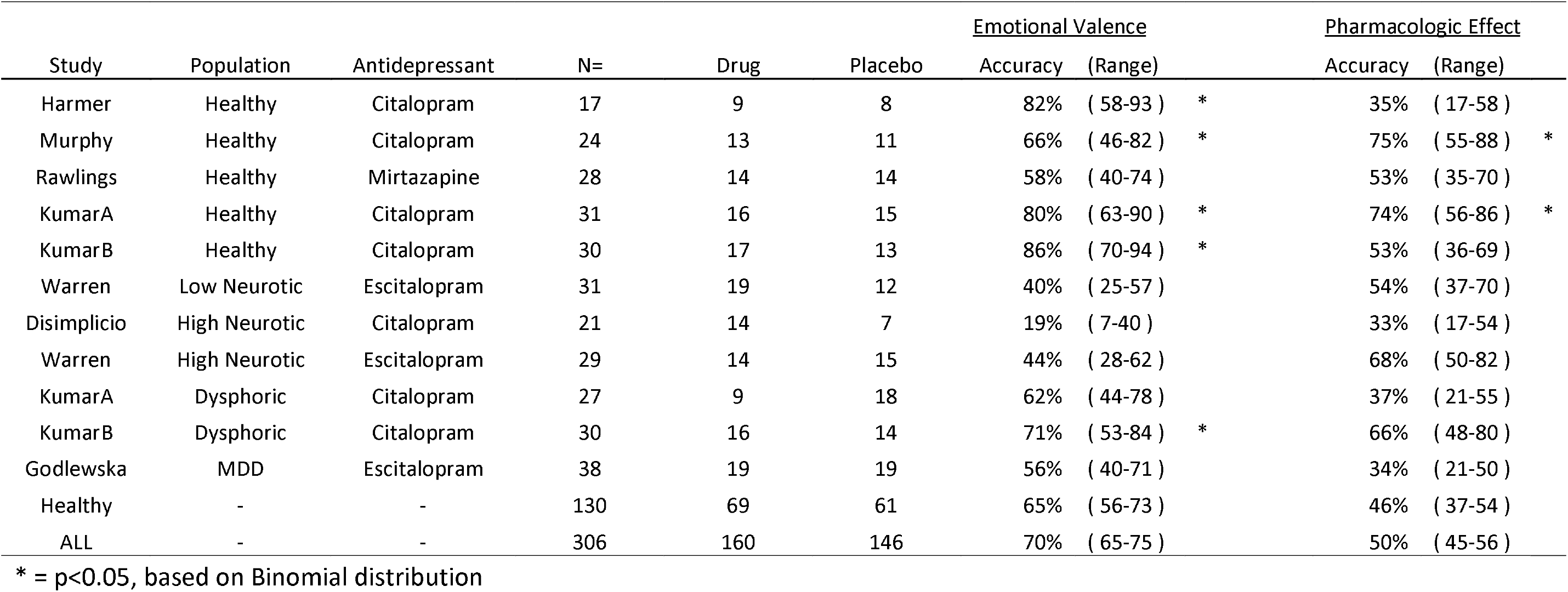
Summary of within-study prediction accuracies. Accuracies give the average (across iterations) proportion of subjects for which the correct contrast was identified. *P* values indicate the probability of achieving this accuracy or better randomly (binomial test, chance = 50%). Range references the Wilson-Score confidence interval (alpha=0.05, sample size as indicated). Shown below are results for the within-study classification and across-all-subjects classification (Healthy, referring only to healthy subjects; ALL, referring to all subjects). Results for the across-study classification may be referenced in Figure 2. Results for the across-all-studies classification may be referenced in Supplementary Figure 2.

In contrast to the above results, we were unable to reliably discriminate the fearful-versus-fixation contrast image from the happy-versus-fixation contrast image better than chance on any classification level (i.e. when fearful and happy were not subtracted within subject, see Supplementary Results).

### 3.2 Pharmacologic Effect: Drug from Placebo Classification

Overall, we found limited ability to identify drug effects in the assessed studies. However, some studies show evidence of a positive drug effect.

i. Within-study classification (i.e. classifiers trained and tested on different subjects of same study) performed best on two of the Kumar datasets representing healthy participants (Kumar A 84% p<0.001; Kumar B *77*%, p=0.01; no test for multiple comparisons as there was one comparison of interest) and dysphoric participants (Kumar A, 74%, p=0.02). These studies also showed a robust effect for emotional valence.
ii. Across-study classification performed poorly. Overall, clinical state had less of an impact on prediction accuracy than the dose and frequency of antidepressant administration. Studies that used 20mg of citalopram or escitalopram for 7 days showed a trend towards higher accuracy than studies that administered a single dose of antidepressant.
iii. Across-all-subjects classification achieved an average accuracy of 56% in held out data. Classifiers trained and tested on only healthy subjects achieved an average accuracy of 50%. These poor results may be associated with the fact that drug and placebo sessions were acquired in separate subjects, so could not be subtracted within subjects, (iv) Across-all-studies classifications achieved an average accuracy no better than chance; these results are presented in Supplementary Results.

## 4. DISCUSSION

We localized an anatomically consistent emotional valence signature in individuals performing the emotional faces task. This valence signature was consistent across subject treatment group (drug or placebo) and drug administration protocol (dose and duration), with similar populations better predicting each other. These results confirm that the emotional valence task strongly probes the brain’s valence circuits notwithstanding differences in task design and clinical population. However, we were unable to find a comparably robust signature for pharmacologic effect. An explanation for this can be inferred from the fact that discrimination of emotional valence was poor when the fearful-versus-fixation contrast image was compared with the happy-versus-fixation contrast image, indicating that the effect of emotional valence could not be isolated from, e.g. te effect of face presentation. As these studies assessed drug effects in parallel groups designs, within-subject drug contrasts were not possible. The present results question the extent to which results from parallel group design studies can generalize using a multivariate machine learning approach. Even minor differences across subjects and across drug protocol are likely to alter measurements of antidepressant effects on the brain’s functional anatomy assessed using this method.

### 4.1 Emotional valence of faces

Our significant classification accuracy for emotional valence suggests that happy and fearful faces engage different aspects of the brain’s functional anatomy in a spatially consistent way across individuals and studies. In this classification, the most relevant parcels included areas reported in meta-analyses of emotional face processing, namely the amygdala and the fusiform gyrus (12). While healthy controls and patients with MDD appear to engage similar functional anatomy during this task, subjects with neurotic traits did not, consistent with previous reports that highly neurotic people have a different response to fear versus happy faces probably as they avert attention and therefore do not process the cues in the same way (13).

In a similar gender-matching emotional faces task, Nord et al. (14) recently reported only moderate (0.4) within-subject, across-trial/day reliability of the BOLD response within the left amygdala and anterior cingulate cortex. Nord et al calculated within-subject reliability for each anatomical area separately, which is perhaps why their results were “surprisingly low.” In our analysis, we evaluated across-subject and across-study reliability in terms of classification accuracy derived from our predictive, multivariate approach which integrated responses from across the entire brain, increasing our sensitivity. Applying a predictive whole-brain approach to investigate within-subject, across-trial reliability will be a useful future analysis.

### 4.2 Pharmacologic effects

Our classifier failed to consistently discriminate drug from placebo. As a general trend, the classifier performed better when trained and tested on similar drug administration protocols, which used the same dose and frequency. And, overall, drug protocols with higher doses for a longer duration (i.e. 20mg for 7 days versus 20mg for 1 day) showed a trend towards higher accuracy. However, we present a guarded interpretation of these results as they could represent false positive results (even though we corrected for multiple comparisons). Why this classification failed could be explained by methodological factors, as well as more general factors that plague drug development studies.

When looking for a subtle signal within the brain’s large-scale networks, individual variability in brain structure and function understandably becomes a significant confounder. Duff et al (2015) reported robust predictions for analgesic studies wherein subjects served as their own placebo control. In addition, each study reported a global effect on brain function that reflected a large pharmacologic effect. Here, we investigated parallel groups-design antidepressant studies where different groups of subjects receiving placebo and drug. While within subject crossover designs could introduce variability associated with order effects, the overall ability to discriminate pharmacologic effects is likely to improve because it will not be muddled with individual variability. Based on our results, we recommend future pharmacologic studies apply a crossover design as a way to minimize individual variability and more ably isolate pharmacologic effect when using classication based machine learning analysis.

In these short, CNS drug administration studies, it is difficult to assess whether therapeutic (here, to affect emotional face processing) CNS drug levels have been reached in each individual due to individual differences in transport proteins that affect blood-brain-barrier permeability (15). Group-wise analyses have shown that acute SSRI administration affects serotonin levels and emotional valence processing in the brain through PET tracer(16) and fMRI studies(17), respectively. Even if the CNS drug levels were known in each individual, however, it would still be difficult to tell whether the same drug level had the same pharmacologic effect in each individual given possible differences in receptor affinity and/or drug coverage. It is further possible that highly localized effects (i.e. like those reported in the largely region-specific antidepressant literature) are diluted and lost in a whole-brain multivariate analysis especially for small areas such as the amygdala which have been consistently reported to be affected with even acute doses of SSRI medication. These factors should be taken into account when interpreting results from CNS-active drug studies.

### 4.3 Implications for Future Work

In the present studies, emotional bias was used to probe the neurobiology of depression based on past group-level observations that depressed individuals have negative emotional bias that corrects with successful treatment (18). While emotional bias is a useful experimental paradigm, the causal connection between emotional bias and depression’s etiology is likely quite complicated. Ramasubbu et al. (19) recently attempted to classify severely depressed patients from healthy controls based on fMRI data alone. They reported statistically significant classification accuracy only for resting-state fMRI data (66%, p=0.012 corrected) while fMRI data acquired during an emotional-face matching task performed at chance. This study suggests that depression may not modulate responses to the emotional-face matching task in a spatially consistent manner. Our results corroborate Ramsubbu et al’s finding by showing that antidepressants seem not to modulate responses to the emotional faces task in a manner that is consistent enough to classify medicated from non-medicated subjects across studies. Because some studies showed some evidence of drug effect, it is possible that a specific, more standardized implementation of the emotional faces task could help probe this antidepressant effect across studies. In comparison to task-based fMRI, it remains to be seen whether resting-state fMRI could more ably probe networks modulated by antidepressants.

This highlights the fact that there are three ontological levels at play in our study: the clinical constructs of healthy, neurotic, dysphoric, and depressed; the experimental construct of emotional bias; and the etiological construct of receptor-specific treatment targets such as the 5-HT receptor. Each represents a different level of analysis. Using symptom-based clinical constructs to probe etiology-based treatments necessarily muddles group treatment effects; similarly diagnosed patients likely have multiple, diverse etiologies. Neurobiology or etiology-based diagnostic categories would likely help isolate the effect of a mechanism-based pharmaceutical by more logically pairing disease etiology with molecular target(20).

Across these levels, it is unlikely that grouping patients by diagnosis (symptom or etiology-based) is the best way forward because patients may have varying symptoms or etiologies within a diagnostic category. Promising research has shown that within a diagnostic category, patients can be grouped by the presence of a specific cluster of symptoms which then predicts their response to a mechanism-based antidepressant (21). This suggests that symptom clusters would serve as reasonable groupings or even features for future predictive analyses. The possibility also remains that a patient’s behavioral performance (in a different experimental construct) will allow a more quantative assessment of a specific cognitive domain, more in line with a dimensional approach to cognitive (dys)function (22).

We suggest that future studies further study the effects of these ontological levels on drug studies and—as much as possible—select more neurobiologically-based means of selecting or probing patient groups.

### 4.4 Methodological considerations

Dimensionality reduction proved a necessary and highly useful step (6). Whole-brain, voxel-wise data (unreported results, wherein each voxel was a feature) were untenable with our available sample size because the number of features greatly outweighted the number of subjects. Whole-brain analysis using data-driven parcellation schemes proved essential in capturing the underlying complex neural circuitry in the brain (10; 23). Given these results, we suggest that future predictive modeling studies use whole-brain parcellation schemes as feature reducers.

An unresolved question is which behavioral task and overall study design best captures the normalizing effect of antidepressants in depression. We evaluated the emotional faces task and discovered differences in effect size which could be based on task presentation, subject population, drug administration protocol or a combination of these. While we report progress in this direction, a more concerted study is required to further address these important questions.

In summary, we applied a cross-validated predictive model to classify emotional valence and pharmacologic effect across eleven task-based fMRI datasets (n=306), exploring the effect of antidepressant administration on emotional face processing. We found patterns of brain activity that successfully classified emotional valence, however could not find such patterns for the pharmacologic effect. Our results also suggest that case-controlled designs and more standardized protocols are required for functional imaging to provide robust biomarkers that can help increase the yield of the drug development pipeline.

## Acknowledgements

This study was funded by Yale University School of Medicine Medical Student ShortTerm Research Funding (D.S.B.) and by the NIH (T32MH019961, R25 MH071584). The authors would like to thank Dr. Kristin S. Budde for her editorial assistance and the staff of Oxford University’s John Radcliffe Hospital’s MRI Centre for their administrative assistance.

## References

1. Insel TR, Sahakian BJ, Voon V, Nye J, Brown V (2012): Drug research: a plan for mental illness. Nature. 483: 269–269.

2. Friedman RA (2013, August 19): A Dry Pipeline for Psychiatric Drugs. The New York Times.

3. Ekman P (2013): Emotion in the Human Face, 2nd ed. (P. Ekman, editor). Los Altos, CA: Malor Books Reprint Edition.

4. Leppänen JM (2006): Emotional information processing in mood disorders: a review of behavioral and neuroimaging findings. Curr Opin Psychiatry. 19: 34–39.

5. Murphy SE, Norbury R, O’Sullivan U, Cowen PJ, Harmer CJ (2009): Effect of a single dose of citalopram on amygdala response to emotional faces. The British Journal of Psychiatry. 194: 535–540.

6. Yoshida K, Shimizu Y, Yoshimoto J, Takamura M, Okada G, Okamoto Y, et al. (2017): Prediction of clinical depression scores and detection of changes in whole-brain using resting-state functional MRI data with partial least squares regression. PLoS ONE. 12: e0179638.

7. Shen X, Tokoglu F, Papademetris X, Constable RT (2013): Groupwise whole-brain parcellation from resting-state fMRI data for network node identification. NeuroImage. 82: 403–415.

8. Jenkinson M, Beckmann CF, Behrens TEJ, Woolrich MW, Smith SM (2012): FSL. NeuroImage. 62: 782–790.

9. Friedman JH (n.d.): Greedy function approximation: a gradient boosting machine. The Annals of Statistics. Vol. 29: 1189–1232.

10. Finn ES, Shen X, Scheinost D, Rosenberg MD, Huang J, Chun MM, et al. (2015): Functional connectome fingerprinting: identifying individuals using patterns of brain connectivity. Nature Publishing Group. 18: 1664–1671.

11. Duff EP, Vennart W, Wise RG, Howard MA, Harris RE, Lee M, et al. (2015): Learning to identify CNS drug action and efficacy using multistudy fMRI data. Sci Transl Med. 7: 274ra16–274ra16.

12. Fusar-Poli P, Placentino A, Carletti F, Allen P, Landi P, Abbamonte M, et al. (2009): Laterality effect on emotional faces processing: ALE meta-analysis of evidence. Neurosci Lett. 452: 262–267.

13. Di Simplicio M, Norbury R, Reinecke A, Harmer CJ (2013): Paradoxical effects of short-term antidepressant treatment in fMRI emotional processing models in volunteers with high neuroticism. Psychol Med. 44: 241–252.

14. Nord CL, Gray A, Charpentier CJ, Robinson OJ, Roiser JP (2017): Unreliability of putative fMRI biomarkers during emotional face processing. NeuroImage. 156: 119–127.

15. O’Brien FE, Dinan TG, Griffin BT, Cryan JF (2012): Interactions between antidepressants and P-glycoprotein at the blood-brain barrier: clinical significance of in vitro and in vivo findings. Br J Pharmacol. 165: 289–312.

16. Nord M, Finnema SJ, Halldin C, Farde L (2013): Effect of a single dose of escitalopram on serotonin concentration in the non-human and human primate brain. Int J Neuropsychopharm. 16: 1577–1586.

17. Rawlings NB, Norbury R, Cowen PJ, Harmer CJ (2010): A single dose of mirtazapine modulates neural responses to emotional faces in healthy people. Psychopharmacology (Berl). 212: 625–634.

18. Harmer CJ, Goodwin GM, Cowen PJ (2009): Why do antidepressants take so long to work? A cognitive neuropsychological model of antidepressant drug action. The British Journal of Psychiatry. 195: 102–108.

19. Ramasubbu R, Brown MRG, Cortese F, Gaxiola I, Goodyear B, Greenshaw AJ, et al. (2016): Accuracy of automated classification of major depressive disorder as a function of symptom severity. NeuroImage: Clinical. 12: 320–331.

20. Barron DS (2016, March 10): Getting Past the “Shotgun” Approach to Treating Mental Illness. Scientific American MIND Guest Blog.

21. Chekroud AM, Gueorguieva R, Krumholz HM, Trivedi MH, Krystal JH, McCarthy G (2017): Reevaluating the Efficacy and Predictability of Antidepressant Treatments. JAMA Psychiatry. 74: 370–9.

22. Insel T, Cuthbert B, Garvey M, Heinssen R, Pine DS, Quinn K, et al. (2010): Research domain criteria (RDoC): toward a new classification framework for research on mental disorders. Am J Psychiatry. 167: 748–751.

23. Rosenberg MD, Finn ES, Scheinost D, Papademetris X, Shen X, Constable RT, Chun MM (2016): A neuromarker of sustained attention from whole-brain functional connectivity. Nature Neuroscience. 19: 165–171.

24. Lundqvist D, Flykt A, Thman A (1998): The Karolinska Directed Emotional Faces-KDEF.

